# Brief Freezing Steps Lead to Robust Immunofluorescence in *Drosophila* Larval Brains

**DOI:** 10.1101/319913

**Authors:** Dominic Buckley, Ada Thapa, Minh Q. Nguyen, Essence Blankinship, Diana Williamson, Veronica Cloud, Ryan D. Mohan

## Abstract

*Drosophila melanogaster* possess complex neuronal networks regulating sophisticated behavioral outputs that aid in studying the molecular mechanisms of neuronal function and neurodegenerative disease. Immunofluorescence (IF) techniques provide a way to visualize the spatiotemporal organization of these networks, permitting observation of their development, functional location, remodeling, and eventually - degradation. However, general immunostaining techniques do not always result in sufficient antibody penetration through the brain, and techniques used to enhance permeability can compromise structural integrity. We have found that freezing larval brains facilitates permeability with no apparent loss of antibody specificity or structural integrity. To demonstrate the advantage of this freezing technique, we compared results to two commonly used permeation methods: Detergent alone (Basic) and proteolytic degradation (Collagenase) techniques.

**Summary:** Here we compare four different immunofluorescence techniques demonstrating that freezing Drosophila brains results in robust staining of small neurons in the larval brain without compromising structural integrity.

## Introduction

*Drosophila melanogaster* is an excellent model organism for neuronal manipulation because it has a relatively moderate number of neurons that command complex behaviors. In addition, a number of neurodegenerative diseases are recapitulated in *Drosophila*, allowing the investigation of changes that occur in neuronal networks due to degenerative processes. One method employed to investigate such changes is immunofluorescence (IF). Various approaches to IF have been developed [1-3], however, efficient permeation of antibodies through the brain can present a problem in producing clear images of target proteins. Permeation using proteolytic enzymes like collagenase [4] can produce more robust staining, but neuronal architecture is disrupted, hindering studies of the spatial organization of neurons. Additionally, proteases can be costly, and other techniques can require day's long antibody incubation [2].

In optimizing IF approaches to examine small groups of cells within larval brains, we found that freezing brains in blocking buffer led to robust staining of target proteins. Importantly, by using freezing methods, brain architecture was preserved. Additionally, freezing time was relatively short, and expensive proteases were not required. Therefore, freezing techniques are useful alternatives to established immunostaining procedures because they allow for clearer visualization of target proteins while avoiding major alterations of brain tissue that occurs in harsher permeation methods. They also save time and money with simple freezing steps.

## Method

To demonstrate the effectiveness of the freezing techniques in immunostaining third instar larval brains, results were compared to those of two other immunofluorescence techniques detailed in the supplemental protocols, basic and collagenase. The protein targeted to show antibody specificity was period, a well characterized neuronal pacemaker protein that is produced specifically in clock neurons [6]. Actin was used to demonstrate staining of large neuronal structures, and nuclei were targeted to show maintenance of brain architecture.

Wild type third instar larval brains (BDSC Cat# 5, RRID:BDSC_5) were dissected in 1X PBS, transferred to a 1.5 ml centrifuge tube, and incubated in 4% formaldehyde for 1 hour at room temperature. Brains were then washed 3 times in 1X PBS by inverting the tube 2 to 3 times. Afterward, brains were washed 3 times in 0.5% PBS Triton X-100 (PBT) by inverting the tube as above. Brains were then incubated in PBT for 20 min at room temperature on a nutator. After removing the PBT from the tube, blocking buffer (5% BSA and 0.2% TWEEN 20 in 1x PBS) was added. Brains were then frozen in solution in one of two ways: The tube was placed at −20°C for 5 min, or brains were frozen by placing the tube on dry ice and spraying the dry ice with 70% ethanol for 10 seconds. After freezing, brains were thawed/blocked in blocking buffer for 1 hour at room temperature on a nutator. Primary anti-period antibody [5] was diluted in blocking buffer to a final antibody dilution of 1:10,000. Blocking buffer was removed from the brains and 200 μl of the primary antibody solution was added. Brains were then incubated in primary antibody at 4°C overnight on a nutator. Afterward, the brains were washed 3 times in PBT by inversion, followed by 4 10-minute washes on a nutator using PBST. Secondary anti-rabbit antibody conjugated to a 594 nm fluorophore (Thermo Fisher Scientific Cat# A-11037, RRID:AB_2534095), and phalloidin labeled with a 488 nm fluorophore (Thermo Fisher Scientific Cat# A12379, RRID:AB_2315147) were diluted in blocking buffer at a 1:1000 and 1:20 dilution respectively. PBT was removed from the tube and 200 μl of the secondary/phalloidin solution was added to the brains. Brains were then incubated in solution for 4 hours at room temperature on a nutator. Afterward, brains were washed 4 times for 10 minutes at room temperature on a nutator using PBT. After removing the PBT, 50 μl of mounting solution containing DAPI (Vector Laboratories, Inc. Burlingame, CA, USA) was added to the brains. Brains were then mounted on microscope slides and allowed at least 30 min to react with the DAPI before imaging on a confocal microscope. Microscope settings were adjusted to capture the best possible image. Settings are presented in Table 1. Representative images of the brain lobes from each technique are shown.

**Table. 1:** WT 3^rd^ instar larval brains were stained using each staining technique: basic, slow freeze, fast freeze, and collagenase. The techniques are detailed in the supplemental protocols. Microscope settings used for each technique are shown in the table above.

|  | Channel | Detector Gain | Amplifier Offset | Pinhole | Wavelength |
| --- | --- | --- | --- | --- | --- |
| Basic | DAPI | 657 | 0.01 | 94 | 405 nm T3 56.6% |
|  | Per | 651 | -0.15 | 136 | 543 nm T1 20.1% |
|  | Phalloidin | 487 | -0.05 | 120 | 488 nm T2 2.1% |
| Slow Freeze | DAPI | 597 | -0.03 | 94 | 405 nm T3 56.6% |
|  | Per | 698 | 0.02 | 124 | 543 nm T1 20.1% |
|  | Phalloidin | 422 | -0.05 | 136 | 488 nm T2 2.1% |
| Fast Freeze | DAPI | 672 | 0 | 94 | 405 nm T3 56.6% |
|  | Per | 721 | -0.13 | 136 | 543 nm T1 20.1% |
|  | Phalloidin | 452 | -0.04 | 120 | 488 nm T2 2.1% |
| Collagenase | DAPI | 702 | 0.01 | 94 | 405 nm T3 56.6% |
|  | Per | 686 | -0.08 | 124 | 543 nm T1 20.1% |
|  | Phalloidin | 452 | 0.01 | 120 | 488 nm T2 2.1% |

## Results

In each technique, period staining is visible in the clock neurons. However, specific staining is most apparent in the collagenase technique, least in the basic technique, and intermediate in the slow and fast freeze techniques. This shows that the collagenase and freezing techniques allow for more robust staining of small neuronal structures deep within the lobes of the brain than basic immunostaining (Fig. 1A). Phalloidin staining reveals stereotypical actin structures for each method. The staining is most apparent in the collagenase technique, followed by the basic, fast freeze, and slow freeze techniques. This demonstrates that the collagenase and basic techniques are better suited for staining larger neuronal structures (Fig. 1B). DAPI staining is similar at the periphery, while staining in the medial portion of the lobes is most apparent in the slow freeze technique, followed by the fast freeze, collagenase, and basic techniques. In addition, the overall shape of the lobes is generally spherical in all but the collagenase technique. In the collagenase method, the lobes come to more of a point at their caudal end. The size of the lobes is enlarged as well, possibly due to disrupted cell junction interactions resulting from the proteolytic activity of collagenase. These results reveal that all but the collagenase technique are best suited for maintaining brain architecture during immunofluorescence (Fig. 1C). Taken together, these results suggest that the slow and fast freezing techniques are preferable alternatives to basic, and proteolytic digestion techniques for staining small structures deep within the brain lobes. The clock neurons are robustly stained in the freezing technique, in contrast to the basic technique, while the integrity of the brain architecture is uncompromised, in contrast to the collagenase technique. However, if staining is to be done on larger neural structures like the actin cytoskeletal network, then the basic technique is the better option since the staining is more apparent than in the freezing techniques, while the structure of the brain is still intact, as opposed to the collagenase technique.

**Fig. 1:**
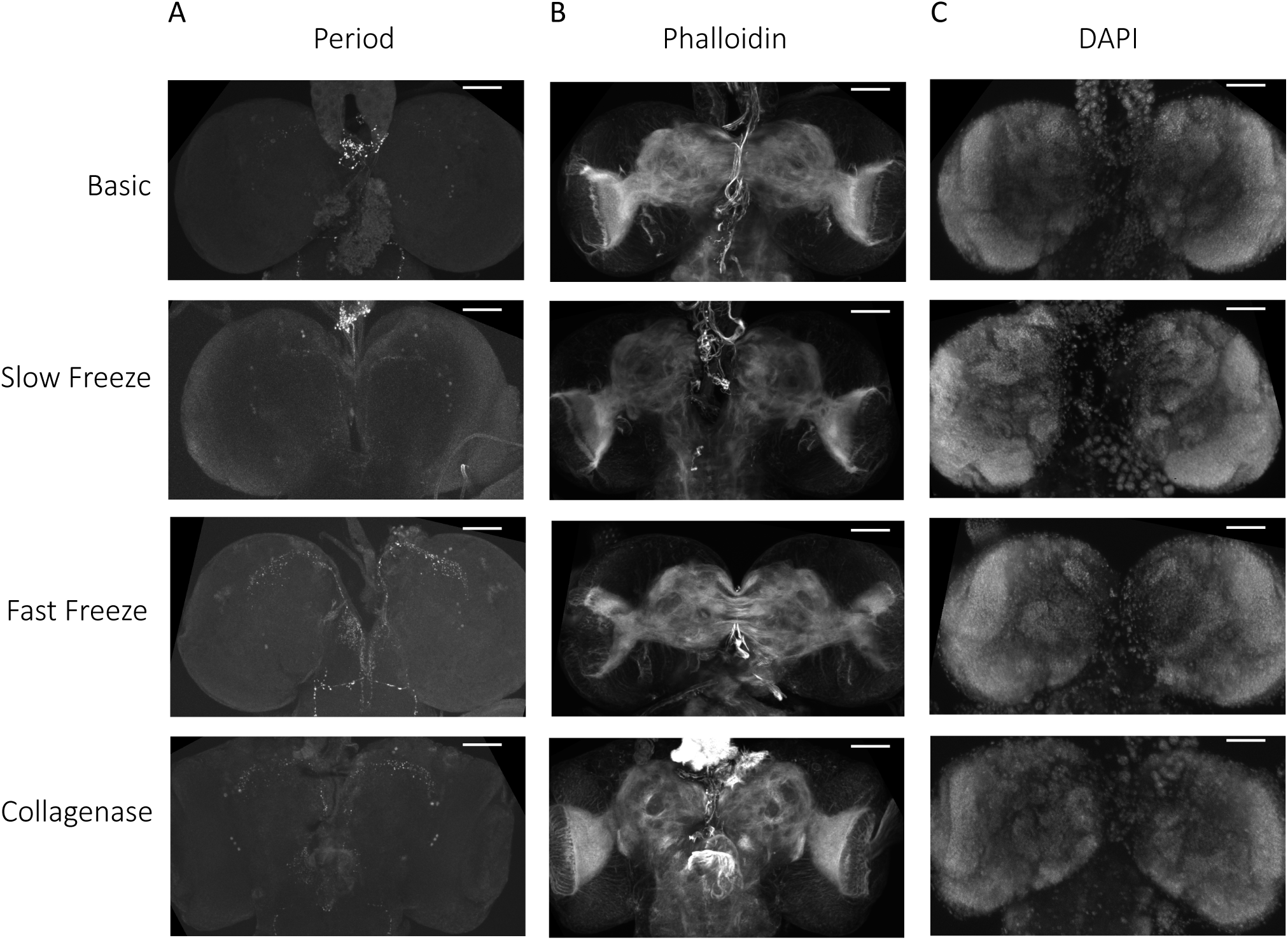
Representative images of 3^rd^ instar larval brain lobes from each immunofluorescence technique. Oregon R 3^rd^ instar larval brains were dissected and stained using each of the four techniques detailed in the protocol. Anti-Per was used to stain clock neurons, phalloidin to stain α-actin, and DAPI to stain nuclei. n=6 brains per technique. Scale bars: 50 μm. Microscope settings were adjusted to capture the best image from each technique. Settings are listed in Table 1.

## Discussion

This benchmark introduces freezing immunofluorescence techniques as alternative methods for obtaining specific antibody staining in Drosophila third instar larval brains without sacrificing sample integrity, time, or money. After optimizing each technique, it is clear that the freezing methods are best suited for staining small neuronal structures deep within the brain lobes, while larger structures in the lobes are best stained using the basic technique. While the collagenase technique does improve staining of neurons, the architecture of the brain is disrupted by the proteolytic treatment, which may disrupt spatial organization of neurons under investigation. Because spatial organization of neurons is important for function, the freezing method is especially useful because the architecture of the brain is not disrupted. This is important for studies of neurodegeneration in *Drosophila* that depend on understanding proper organization of neurons in normal and disease states.

<u>**Protocols**</u>

<u>Basic Immunostaining Technique</u>

## REAGENTS AND MATERIALS

16% Methanol free Formaldehyde solution cat# 15710 (Electron Microscopy Sciences,))
Triton X-100 Cat# T-8787 (Sigma Aldrich, St. Louis, MO, USA)
TWEEN20 Cat# BP337 (Fisher Scientific, Hampton, NH, USA)
Bovine Serum AlbuminA Fraction V Cat# 0332 (VWR Life Sciences, Radnor, PA, USA)
Collagenase (Fisher Scientific, 234153, Hampton, NH, USA)
Primary anti-period antibody (Gift from Jeff and Jin-Yuan Price. See Manuscript)
Secondary anti-rabbit antibody conjugated to a 594 nm fluorophore (Thermo Fisher Scientific Cat# A-11037, RRID:AB_2534095)
Phalloidin labeled with a 488 nm fluorophore (Thermo Fisher Scientific Cat# A12379, RRID:AB_2315147)
Vectashield containing DAPI Cat# H-1500 (Vector Laboratories, Inc. Burlingame, CA, USA)

## PROCEDURE

### Basic Immunostaining technique

Dissection and Fixation of Larval Brains
1. Fill 1 well of a nine well spot plate with 1 ml of 0.03% PBST
2. Fill the rest of the wells with 1 ml of 1x PBS.
3. Place 8 larvae in the 0.03% PBST well and briefly shake each larva using forceps. This is done to wash off any food particles from the larvae.
4. Place each larva into its own well of 1x PBS.
5. Grab the mouth hooks of the larva with one pair of forceps, using the other pair to hold the larva still.
6. Pull the mouth hooks away from the body. The brain lobes and ventral nerve cord are located just caudal to the mouth hooks.
7. After locating the brain lobes and ventral nerve cord, remove all the surrounding tissue.
8. Transfer the dissected larval brains to a 1.5 ml microcentrifuge tube using a p-100 pipette.
9. Remove excess PBS with a p-100 pipette. 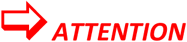 Do not disturb the settled brains for all steps involving the removal of solution from the tube.
10. Add 500 μl of 4% formaldehyde to the tube.
11. Place the tube on a nutator and incubate in formaldehyde for 1 h at room temperature (RT).

Primary Antibody Incubation
11. Remove the tube from the nutator and allow the brains to settle to the bottom of the tube.
12. Remove the formaldehyde from the tube using a P-1000 pipette.
13. Add 500 μl of 1 X PBS to the tube and invert the tube to wash the brains. Allow the brains to settle to the bottom of the tube and remove the PBS. Repeat two times for a total of three washes.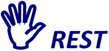 At this point the brains can be stored in PBS at 4°C for up to one month if necessary.
14. Remove the PBS without disturbing the brains.
15. Add 500 μl of 0.5% PBT (0.5% v/v Triton X-100 in 1X PBS) to the tube and invert the tube to wash the brains. Allow the brains to settle to the bottom of the tube and remove the PBT. Repeat two times for a total of three washes.
16. Add 500 μl of PBT to the tube and place the tube on a nutator for 20 min at room temperature (RT).
17. Allow the brains to settle to the bottom of the tube and remove the PBT.
18. Add 500 μl of blocking buffer (5% BSA w/v and 0.2% v/v Tween 20 in 1x PBS) to the tube.
19. Block for 1 h at RT on a nutator.
20. Dilute primary antibodies in blocking buffer. 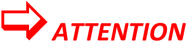 Make at least 200 μl of primary antibody solution and store at 4°C until blocking is finished.
21. Allow the brains to settle to the bottom of the tube and remove the blocking buffer.
22. Add 200 μl of the primary antibody solution to the tube.
23. Incubate the brains at 4°C overnight on a nutator.

Secondary antibody staining
24. Remove the tube from the nutator and allow the brains to settle to the bottom of the tube.
25. Remove the primary antibody solution.
26. Add 500 μl of 0.5% PBT to the tube and invert the tube to wash the brains. Allow the brains to settle to the bottom of the tube and remove the PBT. Repeat two times for a total of three washes.
27. Add 500 μl of PBT to the tube and place the tube on a nutator to wash for 10 min at RT. Allow the brains to settle to the bottom of the tube and remove the PBT. Repeat 3 times for a total of 4 washes.
28. Dilute secondary antibodies 1:1000 in blocking buffer. 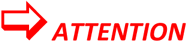 Make at least 200 μl of secondary antibody solution.
29. Remove the tube from the nutator and allow the brains to settle to the bottom of the tube.
30. Remove the PBT from the tube.
31. Add 200 μl of antibody solution to the tube and wrap the tube in aluminum foil. 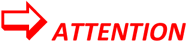 Wrapping in foil is critical because the fluorophores conjugated to secondary antibodies are light sensitive.
32. Place the tube on the nutator at RT and incubate for 4 h.

Mounting Larval Brains
33. Remove the tube from the nutator and allow the brains to settle to the bottom of the tube.
34. Remove the tube from the aluminum foil.
35. Remove the secondary antibody solution.
36. Add 500 μl of 0.5% PBT to the tube and place the tube back in foil. Place the tube on a nutator for 10 min at RT. Allow the brains to settle to the bottom of the tube and remove the PBT. Repeat 3 times for a total of 4 washes.
38. Allow the brains to settle to the bottom of the tube and remove the PBT.
39. Add 50 μl of Vecta Shield with DAPI to the tube.
40. Obtain a microscope slide and two coverslips for mounting.
41. Cut one of the coverslips down the middle with a diamond tip pen and break the cover slip in half along the cut.
42. Place the halves of the coverslips on top of the microscope slide with the straight edges facing each other. Leave a gap between the coverslips.
43. Dab clear fingernail polish on the corners and middle of the outside (jagged) edge of each cover slip to hold them in place.
44. Cut the end of a P-200 pipette tip or use a wide bore P-200 pipette tip to transfer the brains onto the microscope slide between the coverslips. 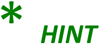 Use forceps to arrange the brains in a straight line down the width of the microscope slide to make finding brains on the confocal microscope easier.
45. Lay the second, intact coverslip down across the cut coverslips to form a “coverslip bridge.” 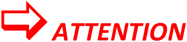 Do not put pressure on the intact cover slip once it has been placed to avoid damaging the brains.
46. Place nail polish along the perimeter of the coverslip bridge to prevent the Vecta Shield from diffusing out.
47. Allow the DAPI to react for at least 30 minutes before imaging using fluorescence microscopy or store the slide at 4°C before imaging.

## PROCEDURE

### Collagenase Technique

Dissection, Collagenase Treatment, and Fixation of Larval Brains
1. Cool 1x PBS on ice.
2. Fill 1 well of a nine well spot plate with 1 ml of 0.03% PBST
3. Fill the rest of the wells with 1 ml of cooled 1x PBS.
4. Place 8 larvae in the 0.03% PBST well and briefly shake each larva using forceps. This is done to wash off any food particles from the larvae.
5. Place each larva into its own well of 1x PBS.
6. Grab the mouth hooks of the larva with one pair of forceps, using another pair to hold the larva still.
7. Pull the mouth hooks away from the body. The brain lobes and ventral nerve cord are located just caudal to the mouth hooks.
8. After locating the brain lobes and ventral nerve cord, remove all the surrounding tissue.
9. Transfer the dissected larval brains to a 1.5 ml microcentrifuge tube using a p-100 pipette.
10. Remove excess PBS with a p-100 pipette. 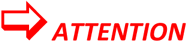 Do not disturb the settled brains for all steps involving the removal of solution from the tube.
11. Add 500 μl of 1x collagenase to the tube and allow to react for 20 min at room temperature (RT).
12. Remove the 1x collagenase from the tube.
13. Add 500 μl of 1x PBS to the tube and invert two to three times to wash away any remaining collagenase. Repeat two times for a total of three washes.
14. Remove the 1x PBS from the tube and add 500 μl of 4% formaldehyde.
15. Place the tube on a nutator and incubate for 1 h at (RT).

Primary Antibody Incubation
16. Remove the tube from the nutator and allow the brains to settle to the bottom of the tube.
17. Remove the formaldehyde from the tube using a P-1000 pipette.18. Add 500 μl of 1x PBS to the tube and invert the tube to wash the brains. Allow the brains to settle to the bottom of the tube and remove the PBS. Repeat two times for a total of three washes.
19. Remove the PBS without disturbing the brains.
20. Add 500 μl of 0.5% PBT (0.5% v/v Triton X-100 in 1xPBS) to the tube and invert the tube to wash the brains. Allow the brains to settle to the bottom of the tube and remove the PBT. Repeat two times for a total of three washes.
21. Add 500 μl of PBT to the tube and place the tube on the nutator for 20 min at room temperature (RT).
22. Allow the brains to settle to the bottom of the tube and remove the PBT.
23. Add 500 μl of blocking buffer (5% BSA w/v and 0.2% v/v Tween 20 in 1x PBS) to the tube.
24. Block for 1 h at RT on the nutator.
25. Dilute primary antibodies in blocking buffer. 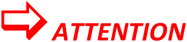 Make at least 200 μl of primary antibody solution and store at 4°C until blocking is finished.
26. Allow the brains to settle to the bottom of the tube and remove the blocking buffer.
27. Add 200 μl of the primary antibody solution to the tube.
28. Incubate the brains at 4°C overnight on a nutator.

Secondary antibody staining
29. Remove the tube from the nutator and allow the brains to settle to the bottom of the tube.
30. Remove the primary antibody solution.
31. Add 500 μl of 0.5% PBT to the tube and invert the tube to wash the brains. Allow the brains to settle to the bottom of the tube and remove the PBT. Repeat two times for a total of three washes.
32. Add 500 μl of PBT to the tube and place the tube on a nutator to wash for 10 min at RT. Allow the brains to settle to the bottom of the tube and remove the PBT. Repeat 3 times for a total of 4 washes.
33. Dilute secondary antibodies 1:1000 in blocking buffer. 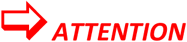 Make at least 200 μl of secondary antibody solution.
34. Remove the tube from the nutator and allow the brains to settle to the bottom of the tube.
35. Remove the PBT from the tube.
36. Add 200 μl of antibody solution to the tube and wrap the tube in aluminum foil. 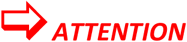 Wrapping in foil is critical because the fluorophores conjugated to secondary antibodies are light sensitive.
37. Place the tube on the nutator at RT and incubate for 4 h.

Mounting Larval Brains
38. Remove the tube from the nutator and allow the brains to settle to the bottom of the tube.
39. Remove the tube from the aluminum foil.
40. Remove the secondary antibody solution.
41. Add 500 μl of 0.5% PBT to the tube and place the tube back in foil. Place the tube on a nutator for 10 min at RT. Allow the brains to settle to the bottom of the tube and remove the PBT. Repeat 3 times for a total of 4 washes.
43. Allow the brains to settle to the bottom of the tube and remove the PBT.
44. Add 50 μl of Vecta Shield with DAPI to the tube.
45. Obtain a microscope slide and two coverslips for mounting.
46. Cut one of the coverslips down the middle with a diamond tip pen and break the cover slip in half along the cut.
47. Place the halves of the coverslips on top of the microscope slide with the straight edges facing each other. Leave a gap between the coverslips.
48. Dab clear fingernail polish on the corners and middle of the outside (jagged) edge of each cover slip to hold them in place.
49. Cut the end of a P-200 pipette tip or use a wide bore P-200 pipette tip to transfer the brains onto the microscope slide between the coverslips. 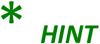 Use forceps to arrange the brains in a straight line down the width of the microscope slide to make finding brains on the confocal microscope easier.
50. Lay the second, intact coverslip down across the cut coverslips to form a “coverslip bridge.” 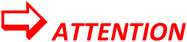 Do not put pressure on the intact cover slip once it has been placed to avoid damaging the brains.
51. Place nail polish along the perimeter of the coverslip bridge to prevent the Vecta Shield from diffusing out.
52. Allow the DAPI to react for at 4 hours at room temperature before imaging using fluorescence microscopy or store the slide at 4°C before imaging.

## PROCEDURE

### Slow Freeze, −20° Celsius Freezing Technique

Dissection and Fixation of Larval Brains
1. Fill 1 well of a nine well spot plate with 1 ml of 0.03% PBST
2. Fill the rest of the wells with 1 ml of 1x PBS.
3. Place 8 larvae in the 0.03% PBST well and briefly shake each larva using forceps. This is done to wash off any food particles from the larvae.
4. Place each larva into its own well of 1x PBS.
5. Grab the mouth hooks of the larva with one pair of forceps, using the other pair to hold the larva still.
6. Pull the mouth hooks away from the body. The brain lobes and ventral nerve cord are located just caudal to the mouth hooks.
7. After locating the brain lobes and ventral nerve cord, remove all the surrounding tissue.
8. Transfer the dissected larval brains to a 1.5 ml microcentrifuge tube using a p-100 pipette.
9. Remove excess PBS with a p-100 pipette. 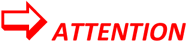 Do not disturb the settled brains for all steps involving the removal of solution from the tube.
10. Add 500 μl of 4% formaldehyde to the tube.
11. Place the tube on a nutator and incubate in formaldehyde for 1 h at room temperature (RT).

Slow Freezing and Primary Antibody Incubation
10. Remove the tube from the nutator and allow the brains to settle to the bottom of the tube.
11. Remove the formaldehyde from the tube using a P-1000 pipette.
12. Add 500 μl of 1 X PBS to the tube and invert the tube to wash the brains. Allow the brains to settle to the bottom of the tube and remove the PBS. Repeat two times for a total of three washes. 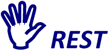 At this point the brains can be stored in PBS at 4° C for up to one month if necessary.
13. Remove the PBS without disturbing the brains.
14. Add 500 μl of 0.5% PBT (0.5% v/v Triton X-100 in 1X PBS) to the tube and invert the tube to wash the brains. Allow the brains to settle to the bottom of the tube and remove the PBT. Repeat two times for a total of three washes.
15. Add 500 μl of PBT to the tube and place the tube on a nutator for 20 min at room temperature (RT).
16. Allow the brains to settle to the bottom of the tube and remove the PBT.
17. Add 500 μl of blocking buffer (5% BSA w/v and 0.2% v/v Tween 20 in 1x PBS) to the tube.
18. Place the tube at −20° C for 5 min.
19. Thaw/block for 1 h at RT on a nutator.
20. Dilute primary antibodies in blocking buffer. 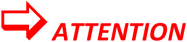 Make at least 200 μl of primary antibody solution and store at 4° C until blocking is finished.
21. Allow the brains to settle to the bottom of the tube and remove the blocking buffer.
22. Add 200 μl of the primary antibody solution to the tube.
23. Incubate the brains at 4° C overnight on a nutator.

Secondary antibody staining
24. Remove the tube from the nutator and allow the brains to settle to the bottom of the tube.
25. Remove the primary antibody solution.
26. Add 500 μl of 0.5% PBT to the tube and invert the tube to wash the brains. Allow the brains to settle to the bottom of the tube and remove the PBT. Repeat two times for a total of three washes.
27. Add 500 μl of PBT to the tube and place the tube on a nutator to wash for 10 min at RT. Allow the brains to settle to the bottom of the tube and remove the PBT. Repeat three times for a total of four washes.
28. Dilute secondary antibodies 1:1000 in blocking buffer. 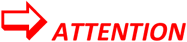 Make at least 200 μl of secondary antibody solution.
29. Remove the tube from the nutator and allow the brains to settle to the bottom of the tube.
30. Remove the PBT from the tube.
31. Add 200 μl of antibody solution to the tube and wrap the tube in aluminum foil. 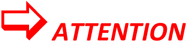 Wrapping in foil is critical because the fluorophores conjugated to secondary antibodies are light sensitive.
32. Place the tube on a nutator at RT and incubate for 4 h.

Mounting Larval Brains
33. Remove the tube from the nutator and allow the brains to settle to the bottom of the tube.
34. Remove the tube from the aluminum foil.
35. Remove the secondary antibody solution.
36. Add 500 μl of 0.5% PBT to the tube and place the tube back in foil. Place the tube on a nutator for 10 min at RT. Allow the brains to settle to the bottom of the tube and remove the PBT. Repeat 3 times for a total of 4 washes.
37. Allow the brains to settle to the bottom of the tube and remove the PBT.
38. Add 50 μl of Vecta Shield with DAPI to the tube.
39. Obtain a microscope slide and two coverslips for mounting.
40. Cut one of the coverslips down the middle with a diamond tip pen and break the cover slip in half along the cut.
41. Place the halves of the coverslips on top of the microscope slide with the straight edges facing each other. Leave a gap between the coverslips.
42. Dab clear fingernail polish on the corners and middle of the outside (jagged) edge of each cover slip to hold them in place.
43. Cut the end of a P-200 pipette tip or use a wide bore P-200 pipette tip to transfer the brains onto the microscope slide between the coverslips. 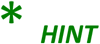 Use forceps to arrange the brains in a straight line down the width of the microscope slide to make finding brains on the confocal microscope easier.
44. Lay the second, intact coverslip down across the cut coverslips to form a “coverslip bridge.” v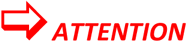 Do not put pressure on the intact cover slip once it has been placed to avoid damaging the brains.
45. Place nail polish along the perimeter of the coverslip bridge to prevent the Vecta Shield from diffusing out.
46. Allow the DAPI to react for 4 hours at room temperature before imaging using fluorescence microscopy or store the slide at 4° C before imaging.

## PROCEDURE

### Fast Freeze Freezing Technique

Dissection and Fixation of Larval Brains
1. Fill 1 well of a nine well spot plate with 1 ml of 0.03% PBST
2. Fill the rest of the wells with 1 ml of 1x PBS.
3. Place 8 larvae in the 0.03% PBST well and briefly shake each larva using forceps. This is done to wash off any food particles from the larvae.
4. Place each larva into its own well of 1x PBS.
5. Grab the mouth hooks of the larva with one pair of forceps, using the other pair to hold the larva still.
6. Pull the mouth hooks away from the body. The brain lobes and ventral nerve cord are located just caudal to the mouth hooks.
7. After locating the brain lobes and ventral nerve cord, remove all the surrounding tissue.
8. Transfer the dissected larval brains to a 1.5 ml microcentrifuge tube using a p-100 pipette.
9. Remove excess PBS with a p-100 pipette. 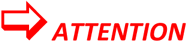 Do not disturb the settled brains for all steps involving the removal of solution from the tube.
10. Add 500 μl of 4% formaldehyde to the tube.
11. Place the tube on a nutator and incubate in formaldehyde for 1 h at room temperature (RT).

Fast Freezing and Primary Antibody Incubation
10. Remove the tube from the nutator and allow the brains to settle to the bottom of the tube.
11. Remove the formaldehyde from the tube using a P-1000 pipette.
12. Add 500 μl of 1 X PBS to the tube and invert the tube to wash the brains. Allow the brains to settle to the bottom of the tube and remove the PBS. Repeat two times for a total of three washes. 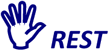 At this point the brains can be stored in PBS at 4° C if necessary.
13. Remove the PBS without disturbing the brains.
14. Add 500 μl of 0.5% PBT (0.5% v/v Triton X-100 in 1X PBS) to the tube and invert the tube to wash the brains. Allow the brains to settle to the bottom of the tube and remove the PBT. Repeat two times for a total of three washes.
15. Add 500 μl of PBT to the tube and place the tube on the nutator for 20 min at room temperature (RT).
16. Allow the brains to settle to the bottom of the tube and remove the PBT.
17. Add 500 μl of blocking buffer (5% w/v BSA and 0.2% v/v Tween 20 in 1x PBS) to the tube.
18. Place the tube in dry ice and spray 70% ethanol around the tube for 10 seconds. 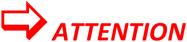 Be sure the solution is frozen. If it is not, spray for a few more seconds.
19. Thaw/block for 1 h at RT on a nutator.
20. Dilute primary antibodies in blocking buffer. 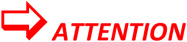 Make at least 200 μl of primary antibody solution and store at 4° C until blocking is finished.
21. Allow the brains to settle to the bottom of the tube and remove the blocking buffer.
22. Add 200 μl of the primary antibody solution to the tube.
23. Place the tube on a nutator and incubate the brains at 4° C overnight.

Secondary antibody staining
24. Remove the tube from the nutator and allow the brains to settle to the bottom of the tube.
25. Remove the primary antibody solution.
26. Add 500 μl of 0.5% PBT to the tube and invert the tube to wash the brains. Allow the brains to settle to the bottom of the tube and remove the PBT. Repeat two times for a total of three washes.
27. Add 500 μl of PBT to the tube and place the tube on a nutator to wash for 10 min at RT. Allow the brains to settle to the bottom of the tube and remove the PBT. Repeat three times for a total of four washes.
28. Dilute secondary antibodies 1:1000 in blocking buffer. 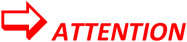 Make at least 200 μl of secondary antibody solution.
29. Remove the tube from the nutator and allow the brains to settle to the bottom of the tube.
30. Remove the PBT from the tube.
31. Add 200 of antibody solution to the tube and wrap the tube in aluminum foil. 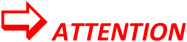 Wrapping in foil is critical because the fluorophores conjugated to secondary antibodies are light sensitive.
32. Place the tube on a nutator at RT and incubate for 4 h.

Mounting Larval Brains
33. Remove the tube from the nutator and allow the brains to settle to the bottom of the tube.
34. Remove the tube from the aluminum foi.
35. Remove the secondary antibody solution.
36. Add 500 μl of 0.5% PBT to the tube and place the tube back in foil. Place the tube on a nutator for 10 min at RT. Allow the brains to settle to the bottom of the tube and remove the PBT. Repeat 3 times for a total of 4 washes.
37. Allow the brains to settle to the bottom of the tube and remove the PBT.
38. Add 50 μl of Vecta Shield with DAPI to the tube.
39. Obtain a microscope slide and two coverslips for mounting.
40. Cut one of the coverslips down the middle with a diamond tip pen and break the cover slip in half along the cut.
41. Place the halves of the coverslips on top of the microscope slide with the straight edges facing each other. Leave a gap between the coverslips.
42. Dab clear fingernail polish on the corners and middle of the outside (jagged) edge of each cover slip to hold them in place.
43. Cut the end of a P-200 pipette tip or use a wide bore P-200 pipette tip to transfer the brains onto the microscope slide between the coverslips. 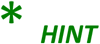 Use forceps to arrange the brains in a straight line down the width of the microscope slide to make finding brains on the confocal microscope easier.
44. Lay the second, intact coverslip down across the cut coverslips to form a “coverslip bridge.” 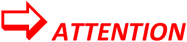 Do not put pressure on the intact cover slip once it has been placed to avoid damaging the brains.
45. Place nail polish along the perimeter of the coverslip bridge to prevent the Vecta Shield from diffusing out.
46. Allow the DAPI to react for 4 hours at room temperature before imaging using fluorescence microscopy or store the slide at 4° C before imaging.

## RECIPES

| 10X PBS | Vendor | Location | amount |  |
| --- | --- | --- | --- | --- |
| NaCl | Sigma Aldrich | St. Louis, MO, USA | 320 (g) | 1.4 M |
| KCL | Thomas Scientific | Swedesboro, NJ, USA | 8 (g) | 26.83mM |
| Na<sub>2</sub>HPO<sub>4</sub> | Fisher Scientific | Hampton, NH, USA | 57.6 (g) | 101.63mM |
| KH<sub>2</sub>PO<sub>4</sub> | Fisher Scientific | Hampton, NH, USA | 9.6 (g) | 17.64mM |
| ddH<sub>2</sub>O |  |  | add to 4 L |  |

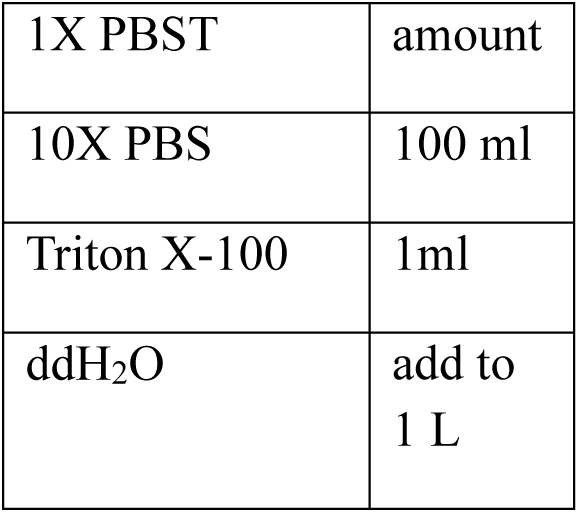

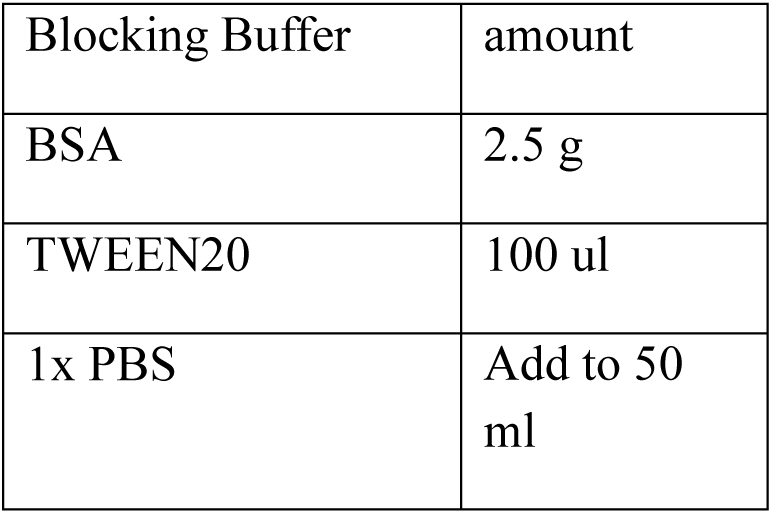

## EQUIPMENT

| Equipment | Vendor | Location |
| --- | --- | --- |
| Leica S6E Stero Microscope | Leica Microsystems | Wetzlar, Germany |
| LSM 5 Pascal Confocal Microscope | Carl Zeiss | Oberkochen, Germany |
| Forceps | Electron Microscopy Sciences | Hatfield, PA, USA |
| Variable Speed Nutator | Fisher Scientific | Hampton, NH, USA |

## TROUBLESHOOTING

Some adjustments may be necessary to optimize this technique. Signal to noise ratio can be improved by adjusting primary and secondary antibody dilutions, time and temperature of incubation in antibody solutions, and time of incubation in blocking buffer. The number of washes and time used to wash the brains can also be adjusted for better signal to noise ratio.

